# Impact of inter-species interactions between flower microbiota commensals and a floral pathogen on disease incidence and pathogen activity

**DOI:** 10.1101/2023.05.16.541000

**Authors:** M. Amine Hassani, Zhouqi Cui, Jacquelyn LaReau, Regan B. Huntley, Blaire Steven, Quan Zeng

## Abstract

Flowers are colonized by a diverse community of microorganisms that alter plant health and interact with floral pathogens. *Erwinia amylovora* is a flower-inhabiting bacterium and a pathogen that infects different plant species, including *Malus × domestica* (apple). Previously, we showed that the co-inoculation of two bacterial strains, *Pseudomonas* and *Pantoea* natural inhabitants of apple flowers, reduced disease incidence caused by this floral pathogen. Here, we decipher the ecological interactions between these two flower-associated bacteria and *E. amylovora* in field experimentation and *in vitro* co-cultures. The two flower microbiota strains did not competitively exclude *E. amylovora* from the stigma habitat, as both commensal bacteria and the pathogen co-existed on the apple flowers, proscribing microbial antibiosis or niche exclusion as predominant mechanisms of host protection. Inspection of the total and the active microbiota populations on flowers revealed that strain co-inoculations altered microbiota activity. Using synthetic stigma exudation medium, tripartite co-culture of the bacterial strains strongly acidified the growth milieu and led to a substantial alteration of gene expression in both the pathogen and the two microbiota members. Our study emphasizes the critical role of emergent properties mediated by inter-species interactions within the plant holobiont and their impact on plant health and pathogen behavior.

**Importance:** Fire blight, caused by *Erwinia amylovora*, is one of the most important plant diseases of pome fruits. Plant-associated microbiota can influence plant disease occurrence through inter-species interactions. Previous studies have shown that plant microbiota commensals could suppress disease mainly by antagonizing of the pathogen growth, however, whether plant-associated microbiota could alter pathogen activity and behavior have not been well studied. Here, we show that the co-inoculation of two bacterial strains that naturally colonize the apple flowers reduce disease pressure. We further demonstrate that the interactions between these two microbiota commensals and the floral pathogen lead to a strong alteration of the pH and to the emergence of new gene expression patterns that may alter the pathogen behavior. Our findings emphasize the critical role of emergent properties mediated by inter-species interactions between plant microbiota and plant pathogens and their impact on plant health.

## Introduction

The flower microbiota provides a useful model system to study the ecological and the evolutionary processes of plant, microbiota and pathogen interactions(1). These short-lived plant organs support the growth of a diverse community of microorganisms(2, 3). The microbial communities that inhabit flowers convey a multitude of functions including interactions with pollinators, altering flower chemistry, and interacting with floral pathogens(1, 4). The inter-species interactions between plant microbiota and plant pathogens may lead to emergent properties that mediate disease resistance(5, 6). Exploring these interactions within the plant holobiont could reveal new mechanisms to control plant disease and promote plant health.

Flowers are nutrient rich microbial habitats that are coveted by plant pathogens(7). The bacterium, *Erwinia amylovora,* is a pathogen that infects several plant species of *Rosaceae* family(8). This bacterium causes fire blight, a disease that incurs significant expenses and losses in pome fruit production. Conventional management of fire blight relies mainly on the prophylactic application of antibiotics, such as streptomycin, oxytetracycline or to a lesser extent oxolinic acid or kasugamycin(9). However, these management strategies favor the evolution of resistance to antibiotics in *E. amylovora* that have been already reported(9). The emergence of antibiotic-resistant strains drives pressure for the development of sustainable practices to control fire blight. Microbial inoculants offer alternative strategies to protect plants against pathogens(10). For examples, the bacterium *Pantoea agglomerans* is known to secrete antibiotics that inhibit *E. amylovora*(11), or *Pantoea vagans* that is proposed to exclude the pathogen from the stigma by competing for limiting substrates(12). Lytic phages have also been shown to provide protection against this disease(13). While the control of the fire blight by these microbial species is mainly mediated through competitive interactions, by direct antagonism or niche exclusion, little consideration is given to the role of emergent properties attribute of inter-species interactions that could mediate host protection(14). Emergent property is defined here by emergence of functions or phenotypes not predictable from the constitute part of the community when observed in isolation(6, 15).

In an earlier study, we showed that the co-inoculation of *Pseudomonas* CT-1059 (*Ps*) and *Pantoea* CT-1039 (*Pa*), two bacterial strains isolated from the flowers of apple, lead to reduced disease incidence in experimental orchards(16). These two microbiota members did not show evidence of strong inhibitory interactions against *E. amylovora* in culture, thus suggesting that ecological interactions among the members likely accounted for the suppression of the disease(16). In the present study, we aimed to decipher the ecological interactions between the two flower-associated bacteria (i.e. *Ps* and *Pa*) and the fire blight bacterium *E. amylovora* using field co-inoculations and laboratory co-culture growth conditions. We hypothesized that bacterial inter-species interactions may lead to emergent properties that alter pathogen behavior and thereby promote plant health. To this end, we investigated the potential of microbial co-inoculation to influence the establishment, survival, and activity of the pathogen *E. amylovora* on flowers. Using synthetic stigma exudation medium(17), we revealed how strain co-cultures altered bacterial growth and gene expression. Through this work, we demonstrated that inter-species interactions lead to emergent properties that were not predictable from binary interactions. We propose that ecological interactions between microbiota commensals and plant pathogens are inherently complex and could lead to emergent functions with significance to plant health.

## Results

### Co-inoculation of two flower bacterial commensals to control fire blight

We have shown previously that the co-inoculation of two flower-associated bacteria, *Pseudomonas* CT-1059 (*Ps*) and *Pantoea* CT-1039 (*Pa*) reduces fire blight occurrence(16). The inoculation of these strains either in combination or as single isolate was repeated in this study. The co-inoculation of *Ps* and *Pa* significantly reduced fire blight incidence to a similar extent as the streptomycin treatment, whereas single isolate inoculation did not significantly reduce the disease incidence (**Fig. 1A**). These results verify our previous findings that co-inoculation of these two flower-associated bacteria reduce fire blight incidence in field experimentation.

**Fig 1.**
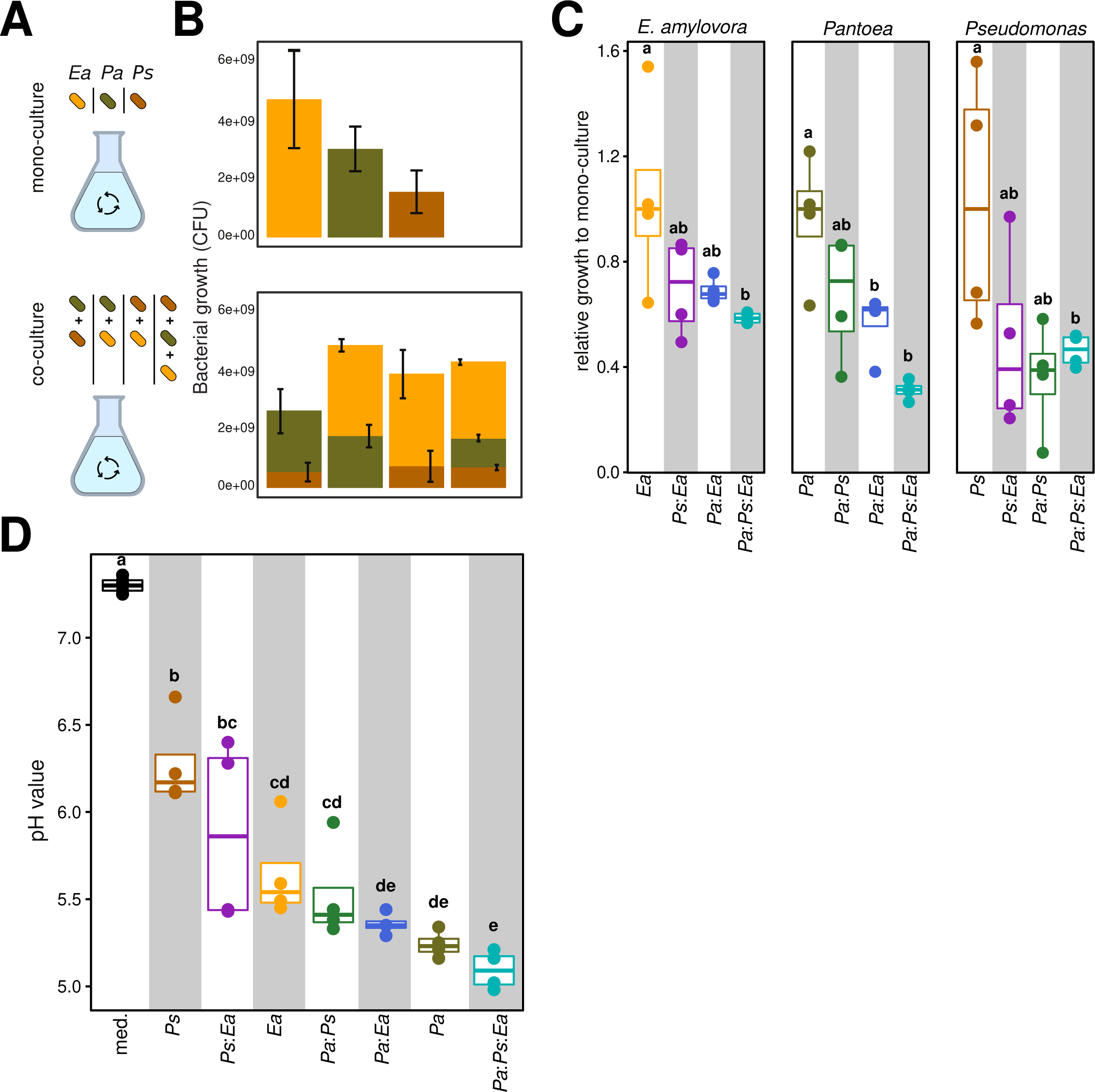
**Co-inoculation of two flower commensal bacteria reduces disease incidence in experimental orchards. A**, the box-plots show fire blight disease incidence in the field under five different treatments: inoculation of *E. amylovora* (*Ea*), inoculation of *Pantoea* and then *Ea* (*Pa:Ea*), inoculation of *Pseudomonas* and then *Ea* (*Ps:Ea*), co-inoculation of *Ps* and *Pa* and then *Ea* (*Pa:Ps:Ea*) or inoculation of *Ea* and then application of streptomycin (*Ea:s*trep.). Co-inoculation of the flowers with both *Pseudomonas* and *Pantoea* significantly reduce disease incidence. Color denotes treatment and letter indicates significant differences between treatments. **B**, the plot depicts *amsC* gene copy number of *E. amylovora* on flower stigma. Each circle corresponds to a single flower and color denotes treatment. *E. amylovora* is detected across all treatments. Neither of the inoculation significantly reduces the population of *Ea* on the flower stigma, except streptomycin. **C**, micrographs show confocal microscopy of *E. amylovora* expressing GFP (top left), *Pseudomonas* expressing mCherry (top right), *Pantoea* expressing mTurqoise (bottom left) and the overlay of the three channels (bottom right) on a flower stigma. *Ps* and *Pa* were pre-inoculation on the stigma, and then *Ea* was applied after 1 day. Microscopy pictures were acquired 2 days post *Ea* inoculation. Microscopy images show that all three bacterial strains are detected and co-exist on stigma habitat.

To investigate whether *Ps* and *Pa* exclude *E. amylovora* 110 (*Ea*) from the flower stigma, we quantified *Ea* abundance by measuring the copy number of *amsC* gene of *Ea* using quantitative PCR (qPCR). The co-inoculation of *Ps* and *Pa* did not significantly alter the absolute abundance of *Ea* on the stigma, whereas we observed that the application of streptomycin did reduce significantly, but marginally, *Ea* abundance (**Fig. 1B**). As qPCR detects DNA, not viable cells, it is plausible that *amsC* genes detected on the stigma originated from dead or inactive *E. amylovora* cells.

To visualize the colonization of the stigma by labeled bacteria that actively express fluorescent proteins (FP), flowers were exposed to *Pseudomonas* CT1181 expressing mCherry FP, *Pantoea* CT-1039 expressing mTurquoise FP, and *E. amylovora* 1189 expressing green FP. Microscopic inspection of the apple flower stigma showed that all three labeled strains were detected and co-occurred on the stigma (**Fig 1C**). Additionally, we observed all three species often co-occurred in close proximity, indicating that even at a micro-scale there was no exclusion of *Ea* in the presence of both probiotic strains. Taken together, these data show that the co-inoculation of *Pa* and *Ps* reduced fire blight disease incidence in the field, but *E. amylovora* is not competitively excluded from the stigma by *Pa* and *Ps*, suggesting other mechanisms mediate plant protection.

### Impact of microbiota structure at different flowering stages on disease incidence

In our previous work, we showed that microbiota structure of apple flowers is temporarily dynamic changing with flower development stages(18). Bacterial communities of apple flowers become stable past the fourth day of flower anthesis(18). Further, the co-inoculation of both bacterial strains (i.e. *Ps* and *Pa*) alters the flower microbiota that resembles the community structure of flowers at late stage of bloom(18). We reasoned that microbial communities of older flowers might be more refractive to the fire blight pathogen and to the disease incidence. To test this, we sprayed *E. amylovora* at Day 2 (young) or Day 4 (older) after anthesis (**Fig 2A**). Flowers were collected at 2 and 4 days post inoculation (dpi) for the samples treated at Day 2 and only at 2 dpi for those treated at Day 4 (**Fig 2A, Methods**).

**Fig 2.**
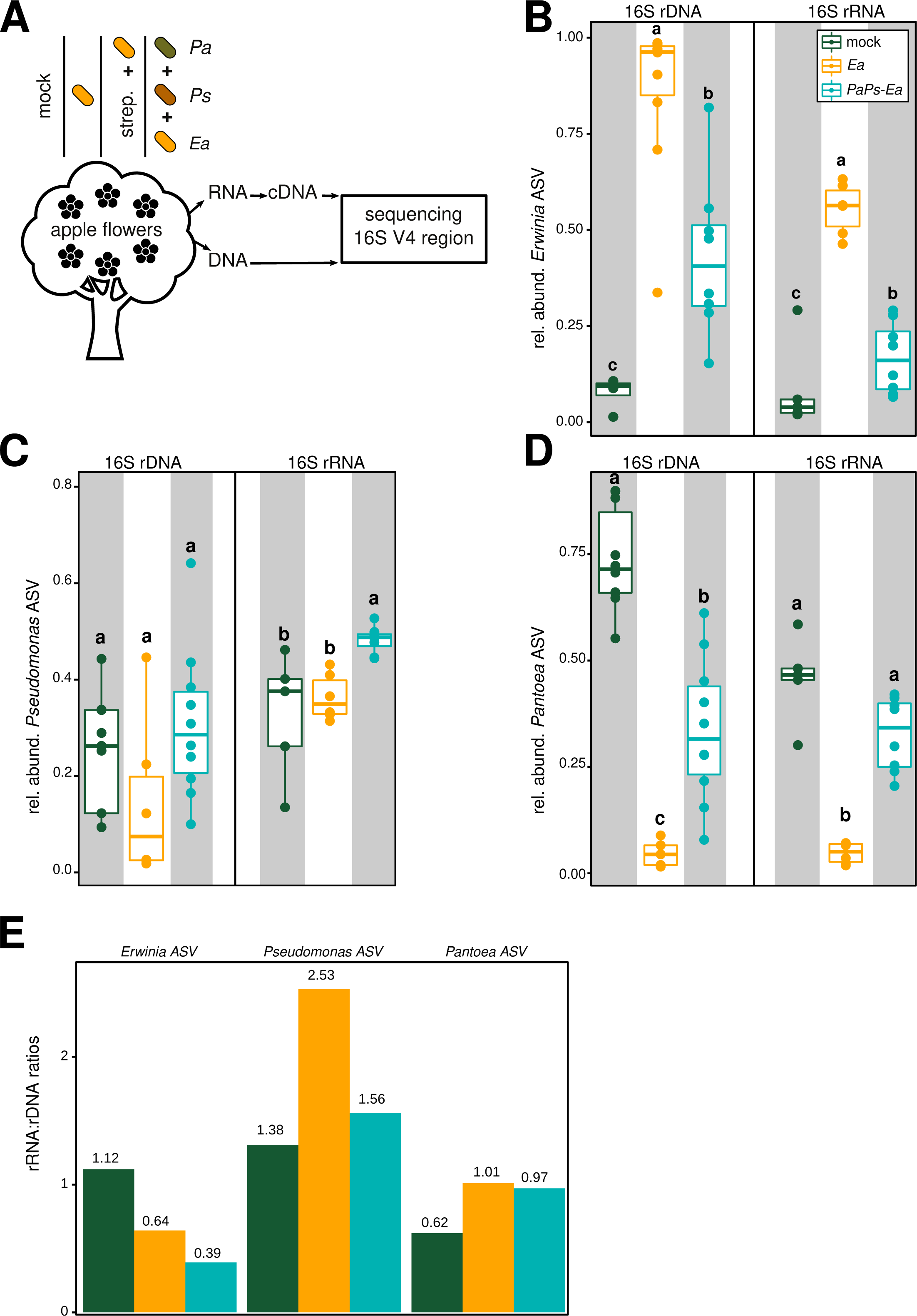
**Inoculation of *E. amylovora* on young and old flowers and its impact on disease incidence. A**, schematic representation that summarizes the experimental procedure. Briefly, flowers were inoculated with *E. amylovora* (*Ea*) at 2 (young) and 4 (old) days after petal opened. Day2-treated flower stigma were harvested at 2 and 4 dpi, whereas Day4-treated stigmas were harvested after 2 dpi. **B**, the box-plots depict copy number of *amsC* gene of *Ea. Ea* reaches higher population density when inoculated at Day 2 than Day 4 (Kruskal-Wallis test, p-value = 0.016, n= 30 flowers). **C**, the box-plots show the apple flower microbiota diversity. **D**, the violin plots indicate Bray-Curtis distances of the apple flower microbiota. R^2^ and p depict explained variance and p-value of the permutational multivariate analysis of variance test, respectively. Inoculation of *Ea* at Day 2 significantly alters the community composition (**C**) and structure (**D**), whereas inoculation at Day 4 does not significantly alter the flower microbiota. Colors indicate either untreated, mock-treated or *Ea*-treated flowers. Each circle indicates the microbiota composition of one individual flower, ten biological replicates were used for each treatment. ** indicate p-value < 0.01 and NS indicate a p-value > 0.05 (Kruskal-Wallis test). **E**, the bar-plots show fire blight disease incidence. Apple flowers inoculated with *Ea* at Day 2 (39.3%) and Day 4 (45.2%) showed comparable disease incidence. Disease was scored 16 days post *Ea* inoculation.

Absolute quantification of *Ea* revealed that higher population densities of the pathogen when inoculated at Day 2 than Day 4 (**Fig 2B**). Although *E. amylovora* reached a smaller population size when sprayed at Day 4, its abundance ranged between 10^6^ to 10^11^ *amsC* gene counts per stigma, suggesting substantial populations of the pathogen could establish on the stigma.

To characterize bacterial communities after the inoculation of *Ea,* we sequenced 16S rRNA genes from the apple flower stigma (**Methods**). By examining observed ASVs (**Fig. 2C**) and Faith’s phylogenetic diversity index (**Suppl. fig. 1A**), we showed that the inoculation of *Ea* at Day 2 significantly reduced bacterial diversity, apparent at both 2 and 4 dpi. In contrast, inoculating *Ea* on Day 4 had no significant effect on bacterial diversity by 2 dpi (**Fig. 2C**). Further inspection of the community structure, via Bray-Curtis dissimilarity, showed that the inoculation of *Ea* at Day 2 induced a shift in the flower microbiota (2dpi: R^2^=0.079, p-value=0.04; 4dpi: R^2^=0.104, p-value=0.002; **Fig. 2D**), whereas *Ea* inoculation at Day 4 did not significantly alter the structure of the microbial community of apple flowers (R^2^=0.054, p-value=0.351; **Fig. 2D, Suppl. fig. 1B**). These analyses reveal that the microbial communities associated with young flowers are less resistant to the disturbance caused by this floral pathogen, whereas those associated with older flowers are not significantly altered by *E. amylovora*. These current results reflect patterns observed in the absolute abundance of *E. amylovora* (**Fig. 2B**), as higher populations of the pathogen altered the diversity and structure of the flower microbiota.

We additionally sought to compare disease incidence between flowers inoculated at Day 2 and Day 4. We found a comparable percentage of symptomatic flowers when inoculated at Day 2 or Day 4 (**Fig. 2E**). Specifically, 39.2% (71/181) and 45.4% (79/174) of the flowers showed blight symptoms (i.e. necrotic wilted flowers) for those inoculated at Day 2 and Day 4, respectively. Collectively, our results indicate that the pathogen *E. amylovora* reaches sufficiently high population densities to cause disease whether sprayed on young (2 day) or older (4 day) flowers. Importantly, these results suggest that the microbiota of developed flowers is not sufficient to suppress fire blight without the co-inoculation of both microbiota commensals.

### Microbial activity in the flower microbiota and the fire blight pathogen

Here, we endeavored to compare the total and the active fractions of the flower microbiota by sequencing 16S rRNA ribosomal genes (rDNA) and 16S rRNA gene transcript (rRNA) respectively(19, 20) (**Methods**), with the expectation that more active populations would be more abundant in the rRNA pools. Apple flowers were either treated with water (mock), with the fire blight pathogen (*Ea*), or with both *Pa* and *Ps* then followed by *Ea* inoculation (*Pa:Ps:Ea*, **Fig. 3A, Methods**). By comparing alpha-diversity measures, we noted that sequencing rRNA captured more bacterial diversity than rDNA in the mock-treated samples (**Suppl. fig. 2A, 2B** and **2C**), suggesting the presence of a diverse active community in the natural microbiota. However, such difference in bacterial diversity between rRNA and rDNA was not observed in microbial-treated samples (**Suppl. fig. 2A, 2B** and **2C**). Profiling the active (rRNA) fraction of the flower microbiota revealed that all microbial treatments were significantly altered in alpha-diversity as compared to mock treatments, whereas total (rDNA) microbiota did not reveal similar community patterns (**Suppl. fig. 2A, 2B, 2C**). Principal coordinate analysis of Bray-Curtis (BC) distances indicated a clear separation between rRNA and rDNA samples (**Suppl. fig. 2D**). Notably, surveying the rRNA revealed a significant shift in the flower microbiota upon the application of *Ea* and the co-inoculation of *Pa* and *Ps* (**Suppl. fig. 2E**: *Ea* and *Pa:Ps:Ea*, respectively). These analyses suggest that the total and the active fractions of the flower microbiota are distinct and profiling the 16S rRNA reveals how the active fraction of the plant microbiota responds to pathogen inoculation.

**Fig 3.**
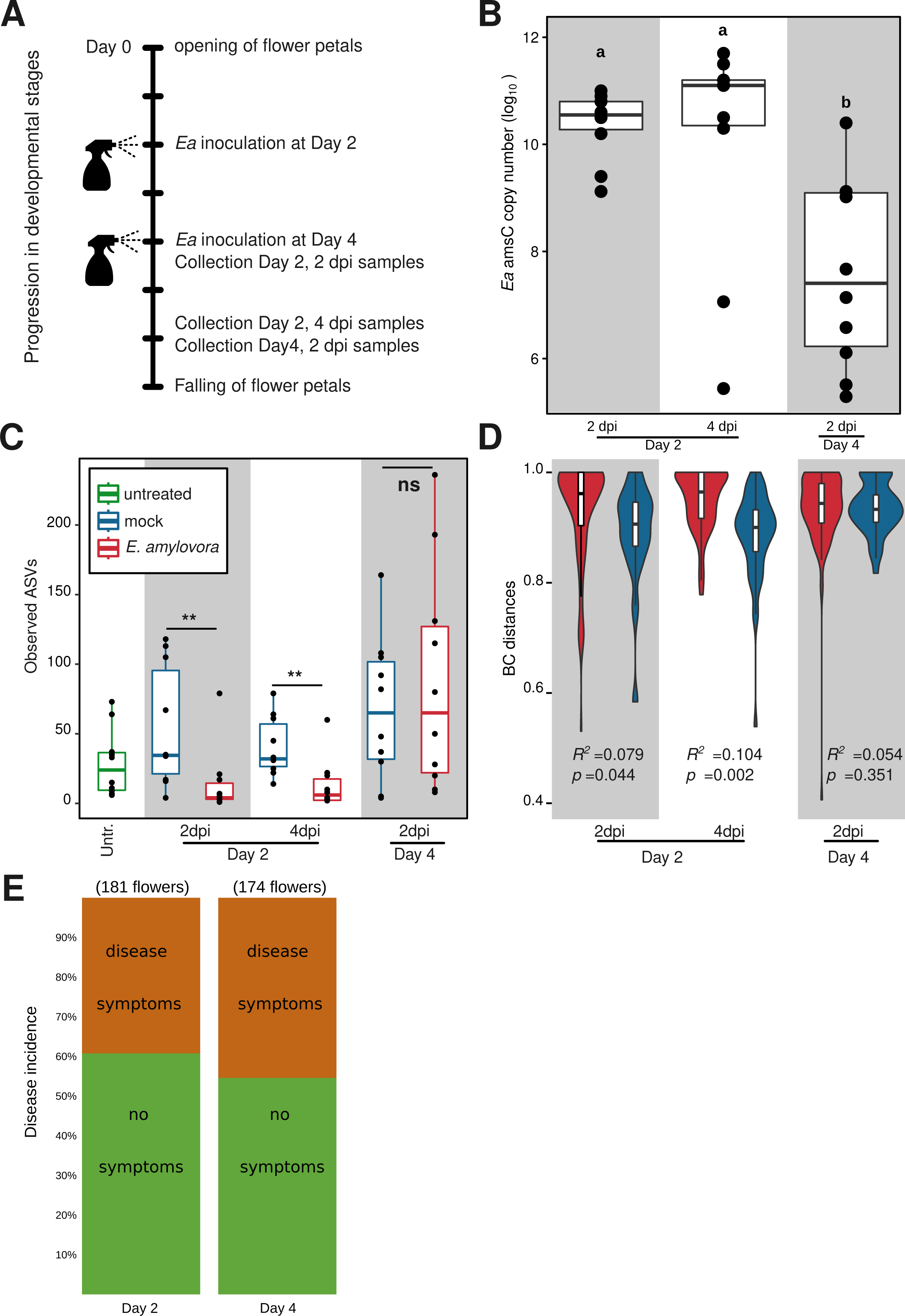
***Pseudomonas* and *Pantoea* alter the activity of *E. amylovora* on the flower stigma**. **A**, the drawing shows the experimental procedure of the co-inoculation of *Pseudomon*as and *Pantoea* on the flowers. Briefly, apple trees were either inoculated with *E. amylovora* (*Ea*), mock-treated or co-inoculated with *Pseudomonas* (*Ps*) and *Pantoea* (*Pa*) during the flowering stage. *Ea* was inoculated to flowers that were pre-inoculated with *Ps* and *Pa*, whereas streptomycin (strep.) was applied to flowers inoculated with *E. amylovra*. Genomic DNA (stigma wash-off from single flower x10 biological replicates) and RNA (stigma wash-off from a pool of 25 flowers x4 biological replicates) were isolated and the V4 region of the 16S gene and transcript were sequenced, respectively. **B, C**, and **D** show the relative abundance of *Erwinia* ASV, *Pseudomonas* ASV and *Pantoea* ASV upon mock treatment, *Ea* inoculation, streptomycin application or *Pseudomonas* and *Pantoea* co-inoculation, respectively. 16S genomic dna and gene transcript show reduction in the relative abundance of *Erwinia* ASV (**B**) and increase in the relative abundance of *Pseudomonas* (**C**) and *Pantoea* (**D**) upon the co-inoculation of *Pseudomonas* and *Pantoea*. **E**, the ratio of the 16S rRNA:16S rDNA relative abundance after the co-inoculation of *Pseudomonas* and *Pantoea* reveals activity of *Ea* compared to *Pa* and *Ps*.

Next, we compared the relative abundance of *Ea, Ps* and *Pa* in rRNA and rDNA sequencing pools (**Fig. 3B, 3C** and **3D**). We noted an increase in the relative abundance of *Ea* in both rRNA and rDNA pools in *Ea-*treated and *Pa:Ps:Ea-*treated samples compared to the mock-treated samples (**Fig. 3B**). These data indicate an increase in both the total and the active populations of the pathogen upon *Ea* inoculation. The relative abundance of *Ps* did not change significantly in the rDNA pool, but it did significantly increase in the rRNA pool upon probiotic co-inoculation (**Fig. 3C**, *Pa:Ps:Ea*), suggesting that *Ps* was more active within the community. The strain *Pa* was more abundant in mock samples as indicated by both rDNA and rRNA pools (**Fig. 3D**, mock). However, *Ea* inoculation led to a substantial drop in the total and the active fractions of *Pa* (**Fig. 3D**, *Ea*). Co-inoculation of bacterial commensals restored the active fraction of *Pa* but remained numerically less abundant on the stigma compared to mock samples (**Fig. 3D**, *Pa:Ps:Ea*). Taken together, these results indicate that bacterial inoculations alter the relative abundance of the total and the active fractions of both commensal bacteria and fire blight pathogen.

It is well established that ribosome copy counts scale with the levels of metabolic activity of the cell(19). Bacterial populations with a higher 16S rRNA:rDNA ratios (>1) show the hallmark of higher cellular activity, whereas lower ratios imply lower levels of cellular activity(20). To test how strain co-inoculations altered the activity of the pathogen and the probiotic bacteria, we computed 16S rRNA:rDNA ratios (**Fig. 3F**). Despite *Ea* having an rRNA:rDNA ratio > 1 in the mock samples (**Fig. 3F**, *Erwinia* ASV, 1.12), it displayed the lowest rRNA:rDNA ratio in conjunction with the co-inoculation of the two flower microbiota strains (**Fig. 3F**, *Erwinia* ASV, 0.39). These findings suggest that co-inoculation of *Pseudomonas* and *Pantoea* (in)directly decrease the activity of the fire blight pathogen. The strain *Ps* displayed rRNA:rDNA ratio of 1.38 in mock samples (**Fig. 3F**, *Pseudomonas* ASV) that further increase to 2.53 upon *Ea* inoculation (**Fig. 3F**, *Pseudomonas* ASV), indicating that *Ps* increased its activity upon pathogen inoculation. The strain *Pa* showed a lower rRNA:rDNA ratio in mock sample (**Fig. 3F**, *Pantoea* ASV, 0.62) that also increased upon *Ea* and probiotic co-inoculation treatments (**Fig. 3F**, *Pantoea* ASV, 1.01 and 0.97, respectively). Thus, *Pa* seems to increase its activity level across the different treatments. Taken together, these analyses show that the active populations of the flower microbiota respond to the pathogen and further indicate that both commensal bacteria alter the activity of *E. amylovora* on the stigma.

### Testing inter-species interaction under co-culture growth conditions

To characterize the inter-species interactions between the two flower commensals and *E. amylovora*, we cultured the bacteria in a synthetic stigma exudation medium(17) either separately (**Fig. 4A:** mono-culture) or in pairwise, or ternary co-cultures (**Fig. 4A:** co-culture). We counted colony forming units (cfu) for each strain by using selective antibiotics (**Methods**). In mono-cultures, the average cell densities of *Ea* reached 4.8 10^9^ cfu/ml, whereas *Pa* and *Ps* reached in average 3.1 10^9^ and 1.6 10^9^ cfu/ml, respectively (**Fig. 4B**: mono-culture). These results show that *Ea* and *Pa* reach higher densities than *Ps* in mono-cultures. In pairwise and ternary co-cultures, total bacterial density reached in average 2.68 10^9^, 3.95 10^9^, 4.37 10^9^ and 4.94 10^9^ cfu/ml in *Pa:Ps, Ps:Ea, Pa:Ps:Ea* and *Pa:Ea* co-cultures, respectively (**Fig 4B:** co-culture). Notably, none of the strains were competitively excluded in any of the pairwise or ternary co-culture conditions (**Fig. 4B:** co-culture). However, we noted that ternary co-culture of the bacterial strains led to significant depletion of *Ea, Pa* and *Ps* by 1.7, 2.2 and 3.0 fold, respectively (**Fig. 4C**). These results show that pairwise and ternary co-culture of *Ea, Pa* and *Ps* using the synthetic stigma exudation medium do not lead to competitive exclusion of any of the test strains, but ternary co-culture leads to a significant growth depletion of all tested bacteria.

**Fig 4.**
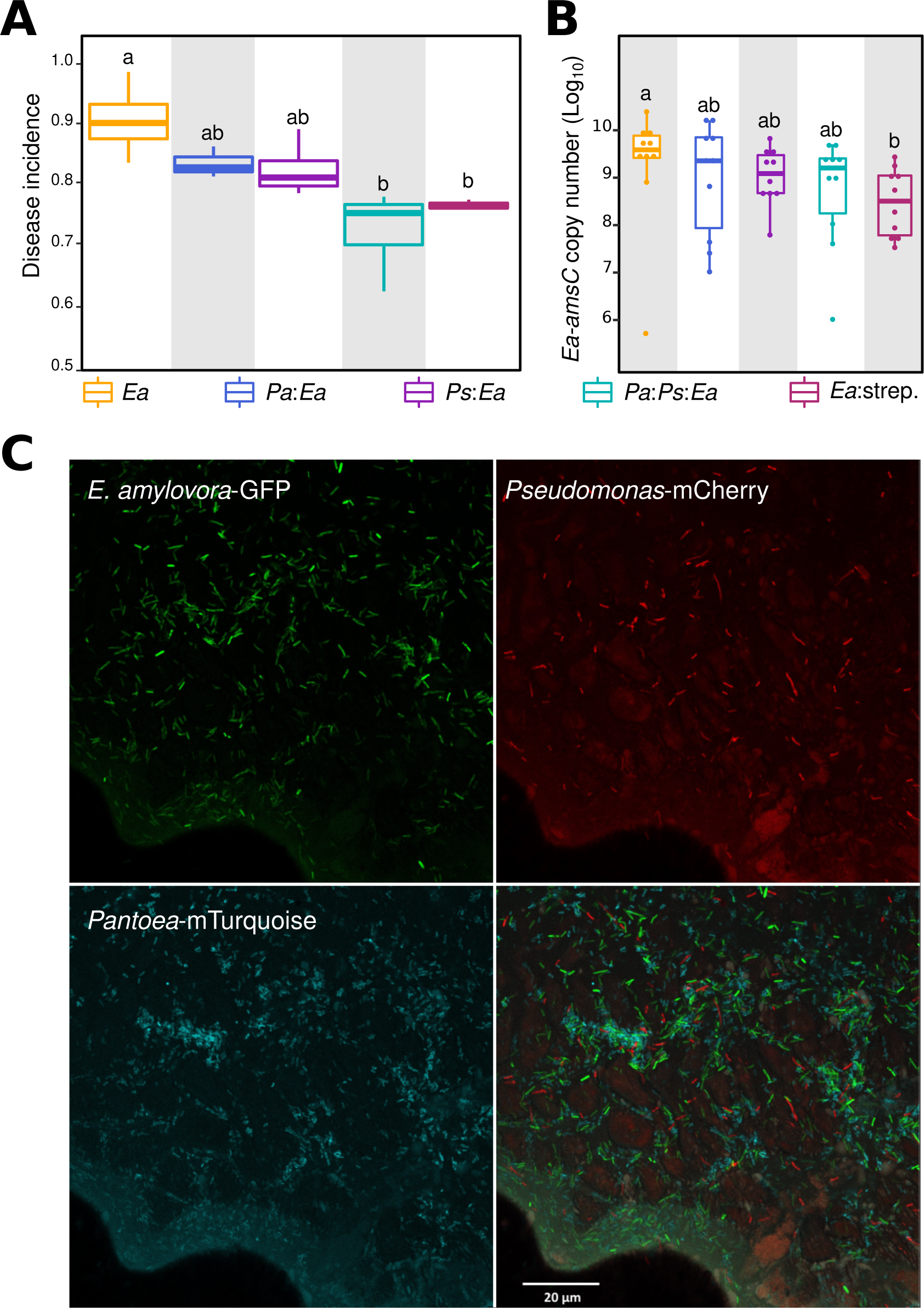
**Tripartite co-culture leads to a strong reduction in the pH of the growth milieu**. **A**, the scheme represents *in vitro* inoculation of *E. amylovora, Pantoea* or *Pseudomonas* **(a),** or combination of their co-inoculations **(b)** into synthetic stigma exudation medium. Bacterial colony forming units (CFU) were determined after 6h post-inoculation using LB agar with a selective antibiotic. The three bacterial species reached different growth rate (top bar-plot panel), but none of them is out-competed (bottom panel). **B**, Upon co-culturing *E. amylovora, Pseudomonas* and *Pantoea*, the bacteria grow significantly, but moderately, less than in mono-culture. C, the box-plots shows pH values in the non-inoculated sigma-mimicking medium and in the mono- and co-cultures. All bacteria significantly alter the pH of the medium, and *Panteoa* strongly shifts the pH of the medium to acidic. Color denotes growth condition and letters show significance in the p-value (p<0.05, Kruskal-Wallis test followe0 d by Conover’s posthoc test).

To test how bacterial inter-species interactions alter the milieu, we measured the pH in each mono- and co-culture growth conditions. We detected pH reduction in all growth conditions (**Fig. 4D**). Mono-culture of *Ps, Ea* and *Pa* reduced the pH on average from 7.30 to 6.27, 5.64 and 5.24, respectively (**Fig. 4D**). The pH of any pairwise co-culture with *Pa* (i.e. *Pa:Ea* and *Pa:Ps*) was more acidic than of the pairwise co-culture of *Ps* and *Ea,* suggesting interactive effects in the final pH of the medium (**Fig. 4D**). Notably, the pH of the ternary co-culture was even more acidic than any pairwise or mono-culture condition (**Fig. 4D**). Thus, while the growth of all three strains alter the pH of the milieu, the ternary co-culture of *Ea, Pa* and *Ps* strongly reduced the pH, supporting interactive effects of these strains in co-culture. Taken together, these findings point to the role of inter-species interactions in altering conditions in the growth milieu that may have consequent properties on microbial interactions and pathogen behavior(21).

### Inter-species interactions modulate the meta-transcriptome

To reveal how gene expression was modulated in bacterial co-cultures, we performed transcriptome analysis for the three strains in different strain assemblages (**Fig. 4A, Methods**). Principal coordinate analysis of gene expression showed clear separation between two groups, one that includes *Ps* and the other that does not include *Ps* (**Fig. 5A**). Thus, the presence of *Peudomonas* strongly induced changes in the meta-transcriptome (**Fig. 4A**).

**Fig 5.**
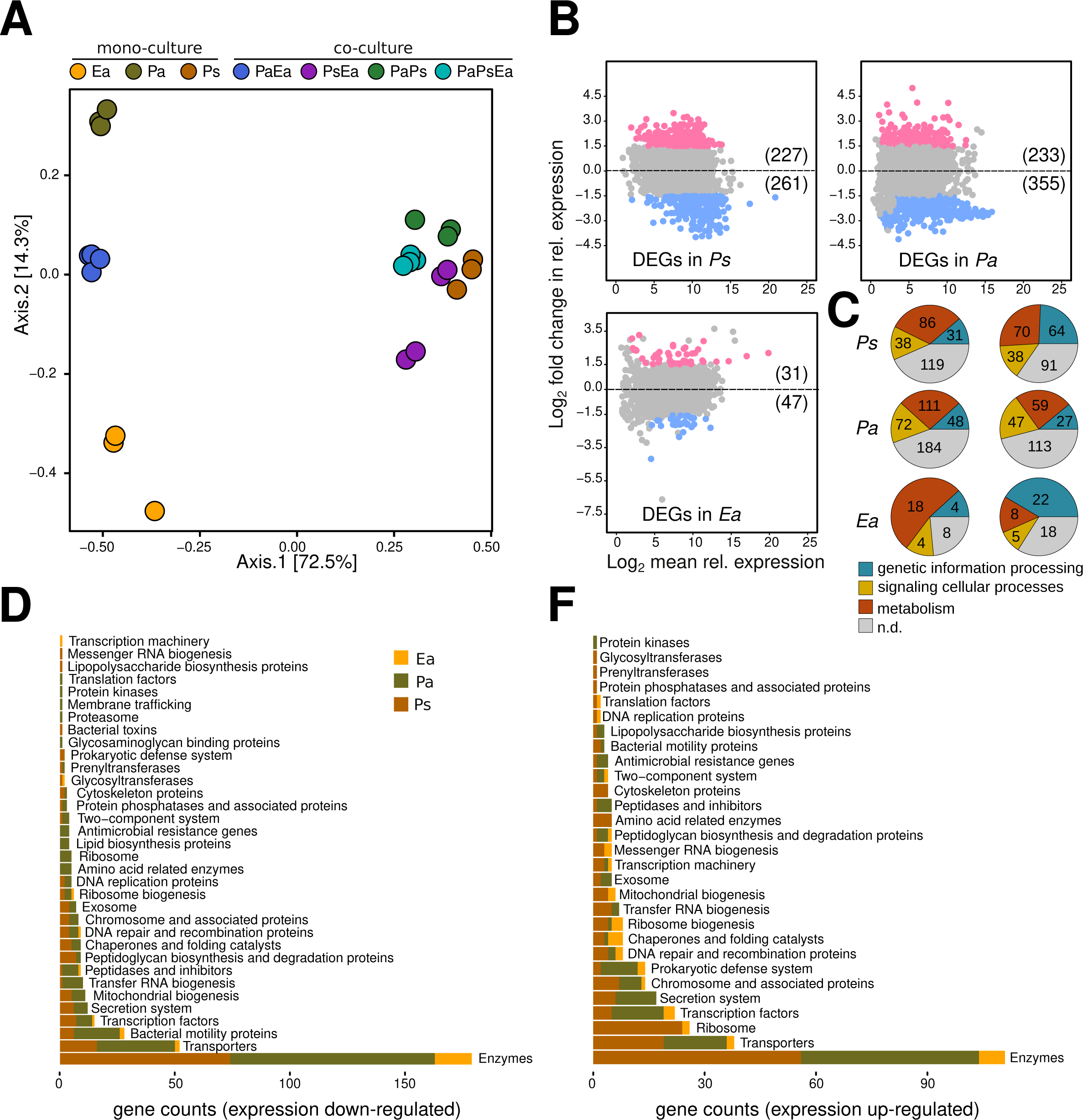
**Meta-transcriptome analysis of bacterial co-cultures. A**, the plot shows principal coordinate analysis in the gene expression profiles of *E. amylovora, Pantoea* and *Pseudomonas* in mono-culture and binary and ternary co-cultures. Bray-Curtis distances between samples were computed on transcript per million of each sample. Color in circles indicates growth condition. *Ea, Pa* and *Ps* show distinct expression profiles and strains co-culture with *Ps* drives the shift in the meta-transcriptome. **B**, top left, top right and bottom left plots indicate differential expressed genes (DEGs) in *Ps, Pa* and *Ea*, upon ternary co-culture, respectively. Count data were fitted to negative binomial distribution (DESeq2) and DEGs were determined based on the cutoffs of 1.5 and 0.05 in log2 fold change expression and Benjamini-Hochberg adjusted p-value, respectively. Red and blue circles indicate up- and down-regulated gene expression upon ternary co-culture. Gray circles indicate genes that were not significantly expressed. Numeric values indicate total number of genes up- or down-regulated in expression. Ternary co-culture of *Pa, Ps* and *Ea* leads to more DEGs in all three strains. **C**, top, middle and bottom charts show proportion of the functional annotation of DEGs in *Ps, Pa* and *Ea*, respectively. Charts in left and right panels indicate DEGs that were down- and up-regulated upon ternary co-culture, respectively. Color code indicates category and n.d. designates not determined. Values inside chart indicate gene counts. **D** and **E** show bar-plots depicting gene ontology using KEGG-BRITE of down- and up-regulated genes, respectively. Color in the bars indicates bacterial strains. The functional category Enzymes counted more DEGs in all *Pa, Ps* and *Ea*.

We computed differentially expressed genes (DEGs) by contrasting relative gene expression in co-cultures *versus* mono-culture growth conditions (**Methods**). The ternary co-culture combination (**Fig. 5B)** led to more DEGs than any pairwise co-culture growth conditions **(Suppl. fig. 3**). For example, we observed 355 (6.51% of the genome) genes with increased expression and 47 (1.35% of the genome) genes with decreased expression in *E. amylovora* grown in a tripartite mixture compared when *Ea* was grown in mono-culture (**Fig. 5B**). In contrast, pairwise co-culture of *Ea* with *Pa* or with *Ps* led to only 8 and 7 DEGs, respectively, compared to the mono-culture of *Ea* (**Suppl. fig. 3C**). Similar patterns were noted for both *Ps and Pa*, with larger numbers of DEGs in ternary cultures than pairwise co-cultures (**Fig. 5B, Suppl. fig. 3**). None of the DEGs in *Ps* and *Ea* were shared across all co-culture growth conditions (**Suppl. fig. 3D** and **3F**, respectively), however we noted 14 down-regulated and 29 up-regulated genes in *Pa* that were differentially regulated across all co-culture growth conditions (**Suppl. fig. 3E:** left and right venn graphs, respectively). These findings highlight that the differentially expressed genes were not additive or predictable from pairwise interactions. Instead, the ternary inter-species interactions led to emergence of new expression patterns for each member of this tripartite community.

To specifically address the genes that were differentially expressed in the ternary growth conditions, the DEGs were functionally annotated. The majority of DEGs in *Ps* and *Pa* were not assigned to a molecular function or a biological process (**Suppl. table 1, Fig. 5C:** n.d. category). The most down-regulated gene category with known functions in all three strains was metabolism, with 31.3% (86 genes), 26.7% (111 genes) and 52% (18 genes) of down-regulated genes in *Ps, Pa* and *Ea* as metabolism-related genes, respectively (**Fig. 5C:** left panel). The genetic information processing and signaling cellular processes were approximately comparable in number of down-regulated genes within strain but not across strains (**Fig. 5C:** left panel). Similarly, metabolism-related genes dominated in percent up-regulated genes in *Ps* (26.6%) and *Pa* (23.9%), except in *Ea* we observed that 41.5% of up-regulated genes were genetic information processing (**Fig. 5C:** right panel). These results show that ternary co-culture leads to the down regulation of metabolism-related genes in *Ea* and indicate shifts in the metabolism of *Pa* and *Ps*. Further classification of DEGs using KEGG-BRITE ontology(22) revealed that most of DEGs in all three strains were of the category Enzymes, with 89, 75 and 16 down-regulated genes (**Fig. 5D**) and 56, 48 and 7 up-regulated genes (**Fig. 5F**) in *Pa, Ps* and *Ea*, respectively. Second most DEGs in all three strains were of the category Transporters (**Fig. 5D** and **5F**). Noteworthy, all three isolates down-regulated genes involved in Bacterial motility (**Fig. 5D**), whereas they up-regulated genes involved in secretion system and prokaryotic defense system when in ternary co-culture (**Fig. 5F**). Altogether, our meta-transcriptomic analysis shows that the three bacterial strains undergo substantial transcriptional re-programming in response to ternary inter-species interactions and highlight the emergence of microbial functions that were not revealed by binary co-cultures.

## Discussion

In this study, we showed that the co-inoculation of two members of the flower microbiota, *Pseudomonas* CT-1059 (*Ps*) and *Pantoea* CT-1039 (*Pa*), reduced fire blight incidence in experimental orchards (**Fig. 1A**). These results corroborate previous findings(16) and further point to the potential use of these strains as probiotic inoculants to control fire blight. Several bacterial strains that control fire blight have been previously identified and used, such as *Pseudomonas fluorescent* A506(23)*, Bacillus subtilis* QST713(24), *Pantoea agglomerans* E325(25), and *Pantoea vagans*(26) to cite few. The mechanism by which above-mentioned bacteria protect the plant from the pathogen is attributed to their ability to reduce or suppress the growth of *E. amylovora* on the stigma through antibiosis or competing for nutrients and space(11, 27, 28). In contrast, our study did not provide evidence of competitive exclusion nor antibiosis of the probiotic strains against *E. amylovora*. Indeed, the abundance of the pathogen remained unchanged on flower stigma and fluorescence micrographs showed that *E. amylovra* co-localize at micro-scale with both flower commensals (**Fig. 1B** and **1C**). Therefore, these bacterial strains do not have adverse effects on the establishment of *E. amylovora* populations, but rather point to the ecological coexistence of the three bacteria on the flowers. However, the co-inoculation of *Pseudomonas* and *Pantoea* strains was required to maximize plant protection. This was corroborated by the observation that both flower cohorts treated at Day 2 and Day 4 developed blight symptoms at similar rates (**Fig. 2E**). Thus, the observed reduction in disease incidence was not due to the microbiota structure but rather point to the role of inter-species interactions leading to emergent properties that may modulate pathogen behavior.

Because the commensal bacteria (i.e. *Pa* and *Ps*) did not impede the colonization of *Ea*, we sought to investigate how these strains influence the activity of the pathogen. We first tested the activity of the flower-associated strains and the pathogen by computing 16S rRNA:rDNA ratios(20, 29). While it has been pointed out the potential pitfalls in 16S rRNA:rDNA ratio analyses(30), these analyses hold suitable to identify the most active populations in a microbial community(31, 32). Our results show that all bacterial strains responded to the treatments by altering their activity and that the co-inoculation of *Pa* and *Ps* reduced the relative activity of *E. amylovora*. It remains unclear whether alteration of the pathogen activity is mediated by direct microbe-microbe interactions or indirectly *via* alternation of the growth conditions on flowers.

To reveal how the two commensal bacteria could alter the activity of the pathogen, we used a growth medium that partially reconstitute components of stigma exudates(17) and revealed how binary and ternary interactions between *Pa, Ps* and *Ea* may alter their growth, pH and gene expression. None of the tested strains were competitively excluded in any of the co-culture conditions (**Fig. 4C**), suggesting the co-existence of these strains in the tested growth medium. Notably, ternary co-culture of *Pa, Ps* and *Ea* led to a strong reduction in the pH of the milieu (**Fig. 4B**) and depleted the growth of all three strains (**Fig. 4C**). The strain *Pa* was the strongest driver of pH reduction. Pusey and colleagues have shown that *Pantoea agglomerans* strain E325 acidifies the pH on the flowers to a point that could deplete the growth of the fire blight pathogen. However, the authors also demonstrated that *P. agglomerans* E325 exhibits antibiosis activity against *E. amylovora*(25, 33). Our study did not provide evidence for antibiosis activity of *Pantoea* against *E. amylovora*, which points to alternative mechanisms of host protection. Further, Pester and colleagues have shown that reduction of pH on apple flowers lead to the down expression of virulence genes in *E. amylovora*(34). Similar dynamics have been described for other pathogen-host systems. For instance, the bumble bee symbiont, *Lactobacillus bombicola*, was shown to inhibit the pathogen *Crithidia bombi* by reducing the pH in the gut(14). These studies, highlight the role of altering environmental pH as a potential property that modulates microbial interactions and pathogen behavior(21, 35). Whether the probiotic strains acidify the stigma and to what extent this may influence the pathogen virulence remain to be solved.

The meta-transcrtiptome study revealed that tripartite interactions between *Pa, Ps* and *Ea* led to a re-programming of gene expression in the pathogen and both commensal bacteria (**Fig. 5A**). A hallmark of this shift was the substantial number of differentially expressed genes (DEGs) in the three strains upon ternary interactions (**Fig. 5B** and **Suppl. fig. 3**). Characterization of DEGs (**Suppl. table 1**) revealed that several microbial functions were associated with the shift in the meta-transcriptome, including acidic stress response, metabolism and microbe-microbe interaction (**Suppl. fig 4A, 4B** and **4C**). For instance, we noted the down expression of genes involved in the metabolism of pyruvate in *E. amylovora* when in co-culture with *Pantoea* and *Pseudomonas* (**Suppl. Table 1**, Ea_1294). Notably, pyruvate was shown to be involved in acid resistance mechanisms in bacteria(36). Also, we observed up-regulation of *rcsB,* a two-component signal transduction system that controls acid resistance in bacteria(37). Another example is illustrated by the up-regulation in the expression of spermidine export protein, an important molecule for cell growth under acidic condition(38)(**Suppl. Table 1:** Ea_02894). These findings indicate that growth conditions under ternary co-culture lead to acidic stress response in the pathogen and further suggest that alteration of the pH may also contribute to the transcriptional re-programming in the bacteria. Re-programming of gene expression was not limited to the pathogen, ternary co-culture of the three strains led to substantial alteration in the transcription patterns of both flower commensal bacteria (**Fig. 5B**). The expression of several genes that encode for ABC transporters were either up- or down-regulated in *Pseudomonas* (**Suppl. fig 4A:** *Ps*) and *Pantoea* (**Suppl. fig 4A:** *Pa*), suggesting that these bacteria undergo shifts in metabolism. Also, we noted alteration in the expression of genes that mediate (in)direct bacterial inter-species interactions. Under ternary co-culture condition, *Ps* up-regulated the expression of a gene that encode for pyroverdine, an iron chelating molecule(39). This siderophore molecule could be employed by bacteria to impede the growth of competitor strains(40, 41). The increase in the expression of genes that encode of type VI secretion in *Pseudomonas* and conjugal transfer pilus in *Pantoea* added further evidence of bacterial inter-species interactions in the tripartite co-culture (**Suppl. fig. 4C**). It is well known that the type VI secretion system mediate direct bacterial killing by injecting toxic proteins into competitor cells(42). Whereas the conjugal transfer pillus has been proposed to mediate secretion of toxins(43). Interestingly, the three bacterial strains up-regulated genes involved in prokaryotic defense system, such as toxin-antitoxin (TA) systems (**Suppl. Fig. 4C**). TA systems have been shown to mediate functions such as persistence, protection against mobile genetic elements and phage inhibition(44). As bacteria could use phages as competitive mechanism against their opponent microbes(45–47), it is plausible that these molecular systems are used as defense mechanisms. Taken together, the meta-transcriptomic study revealed that both flower-associated bacteria and the floral pathogen undergo substantial re-programming in gene expression and the emergence of microbial functions non-additive of the binary co-cultures.

In conclusion, this study showed that the co-inoculation of two flower-associated bacteria protect the host plant from the fire blight disease. These two bacterial strains did not competitively exclude *E. amylovora* from the stigma habitat, but bacterial inter-species interactions altered strains activity. Ternary interactions between the bacteria led to a strong alteration of the pH and to the emergence of new expression patterns. Our study emphasizes the role of emergent microbial properties that are not predictable from binary interactions. We propose to further explore inter-species interactions between microbial consortia and plant pathogens to mediate disease resistance. As the outcome of inter-species interactions are inherently complex, we caution that microbial virulence is an emergent microbial propriety(48), thus deciphering these interactions requisite empirical testing. Exploring the properties of microbiota commensals offers alternative and sustainable practice to manage plant diseases and promote host health.

## Methods

### Bacterial strains and growth conditions

Bacterial strains used in this study were *Erwinia amylovora* strain 110, *Pseudomonas* CT-1059 (field co-inoculations and *in vitro* co-culture assay) and *Pantoea* CT-1039 (field co-inoculations, *in vitro* co-culture assay and microscopy imaging), *Pseudomonas* CT1181 (microscopy imaging). All strains were cultured in Lysogeny Broth (LB) medium at 28°C overnight with shaking (250 rpm). *E. amylovora* and *Pseudomonas* were naturally resistant to rifampicin and streptomycin, respectively. *Pantoea* was sensitive to both aforementioned antibiotics. Antibiotics were added at the following concentrations when required: chloramphenicol, 30 µg/mL; kanamycin, 50 µg/mL; ampicillin, 100 µg/mL; spectinomycin, 50 µg/mL.

### Bacterial strain co-inoculations in the field and fire blight scoring

Field co-inoculations were performed on 30-year-old apple tree cultivar ‘Red Delicious’ in May 2021 at the Lockwood Farm, Hamden, CT (41.406 N, 72.906 W). 36 apple trees were randomly grouped into nine treatment groups with four tree replicates per treatment and organized in a randomized complete block design. Treatments include 1. water, 2. *E. amylovora* 110 (*Ea*), 3. *Pantoea* sp. CT-1039 (*Pa*), 4. *Pseudomonas* sp. CT-1059 (*Ps*), 5. *Pa-Ps,* 6. *Pa:Ea,* 7. *Ps:Ea,* 8. *Pa:Ps:Ea,* and 9. streptomycin. *Pantoea* sp. CT-1039 and *Pseudomonas* sp. CT-1059 were cultured in Lysogeny Broth (LB) overnight. Cells were pelleted by centrifugation at 5,000 rpm for 15 minutes, adjusted to ca. 5 x10^7^ CFU/ml in water, and then sprayed to apple flowers approximately 2 liters per tree. Treatments were applied twice, at 60% bloom (April 29^th^, 2021) and at 80% bloom (April 30^th^, 2021), using a motorized Solo sprayer (3.8 L per tree). *E. amylovora* was inoculated at 100% bloom (May 1^st,^, 2021) with the concentration of 5 x 10^6^ CFU/ml. Streptomycin (Firewall 50, at 100 ppm) was applied 2 hours post *Ea*-inoculation (May 1^st,^, 2021). For *Pa:Ps* co-inoculation, the bacteria were mixed at equal volume and applied to same dosage as single strain treatments. Fire blight disease infection was rated 3 weeks after the inoculation of *E. amylovora*. Disease incidence was calculated as percentage of symptomatic flower clusters in total flower clusters of each tree per treatment.

### Flower sampling for *E. am*ylovora quantification and community profiling

From the field co-inoculation experiments conducted May 2021, we collected flower samples for DNA and RNA extractions. DNA from the sampled flowers were used to quantify *E. amylovora*, and to profile microbial communities using 16S rRNA gene, whereas RNA from the sampled flowers were used to profile the flower microbiota using 16S rRNA gene transcripts. Flowers were harvested after 2 days post-inoculation of *E. amylovora* (dpi, May 3^rd^). For DNA isolation, 10 individual flowers for each treatment were harvested. Four to five stigmas from each individual flower were dissected, placed into micro-centrifuge tube using sterile scissors, snap frozen in liquid nitrogen and were treated as a single biological replicate. For RNA extraction, stigmas were collected from 20 flowers, placed into micro-centrifuge tube. snap frozen and treated as one biological replicates. Samples were stored at -80°C till further processing.

### Absolute quantification of *E. amylovora* by qPCR

The absolute abundance of *E. amylovora* in each stigma sample was quantified using a previously described method(49). In brief, DNA copy number of *E. amylovora* was quantified by determining the cycle threshold (CT) value of the *E. amylovora* specific gene *amsC*. Quantitative PCR (qPCR) was performed using SsoAdvanced universal SYBR Green supermix (Bio-Rad, CA, USA) on a Bio-rad CFX96 realtime PCR, as described previously(18). The CT values for a 1/10 dilution series of known *amsC* gene copies of *E amylovora* chromosomal DNA was determined to make a standard curve for calculation of copy numbers in stigma samples.

### Confocal microscopy observation of microbial colonization on stigma

*Pseudomonas* CT-1181, *Pantoea* CT-1039 constitutively expressing mCherry (red) and mTurqoise (turqoise) fluorescent proteins (FP) were generated using miniTn7-pURR25DK-3xmCherry and miniTn7-pURR25DK-mTurquoise, respectively(50). Using tri-parental mating including helper strain *E. coli* RHO3 carrying pTNS, mCherry and mTurqoise were insert at unique site *att*Tn7 in the genome of recipient *Pseudomonas* CT-1181 and *Pantoea* CT-1093, respectively(51–53). Derivative *E. amylovora* 1189 expressing green fluorescent protein carried a *gfp* (green) under the control of *nptII* promoter integrated into the chromosome. Bacterial cells were cultured in LB overnight at 28°C shaking conditions (250 rpm). Cells were pelleted by centrifugation and adjusted to 10^7^ CFU/ml before they were spray-inoculated onto the stigma of 3 years old trees ‘Gala’ individually potted. Inoculated trees were kept in a plant growth chamber (25° C, 80% relative humidity and 16-hour cycle of 350 µmol light intensity). *Pseudomonas* CT-1181 expressing mCherry FP *and Patoea* CT-1039 expressing mTurqoise FP were inoculated on stigma when petals first open, *E. amylovora* 1189 expressing green FP was inoculated 1 day thereafter. Stigmas were collected 2 days post the inoculation of *E. amylovora*. Fluorescent green (484 nm/507 nm), mCherry (587 nm/610 nm) and mTurqoise (434 nm/474 nm) signals were observed with a Leica TCS SP5 confocal microscope (Leica Microsystems, Wetzlar, Germany) equipped with four laser channels (405 nm, multiline Argon, 561 nm and 633 nm) and two HyD detectors. Images were captured with Leica LAS AF software and overlayed using Leica LAS-X software.

### Field experiment of *E. amylovora* inoculation at different flowering stages

These experiments were performed on 22-year-old apple cultivar ‘Early Macoun’ (4-NY75414-1) in May 2021 at Lockwood Farm, Hamden, CT (41.406 N, 72.906 W). Eight trees were used to tag flower clusters that opened at the same time. Flowers were inoculated with *E. amylovora* 110 at Day 2 or Day 4 after the petals opened. *E. amylovora* 110 was adjusted to 10^7^ CFU/ml in sterile water, and then sprayed to open apple flowers using a hand-held sprayer. A mock treatment of sterile water was also sprayed to open flowers at the same time. Treated flowers were collected 2 and 4 days post inoculation (dpi) for Day 2 and at 2 dpi for Day 4 treatments, respectively. Stigmas from one individual flower were dissected, transferred into a 1.5 ml sterile tube, snap frozen in liquid nitrogen and then stored at -80° C till further DNA processing, and were considered as one biological replicates. Detailed protocol for DNA extraction is indicated further below. Flowers used for quantifying disease incidence were left on the tree for an additional 16 days prior to evaluation of fire blight symptoms. Fire blight symptoms scoring were evaluated as described above.

### DNA extraction from apple stigma

Deep frozen stigma samples were retrieved from -80 °C and 200 µl of 1X PBS (pH 7.4, Invitrogen, Carlsbad CA) + 0.001 % Silvet L77 (PlantMedia, Dublin OH) were added to the tube. To retrieve epiphytic bacteria from stigma surface, samples were sonicated for 5 min. in a water batch at room temperature followed by full speed vortex for 30 sec. Stigma tissues were removed from tube and DNA was extracted from the solution using DNeasy PowerSoil Pro Kit (Qiagen, Hilden Germany) following manufacturer’s instructions except the following modifications. Cell suspensions were treated with 36 µl of Lysozyme (10 mg/ml in PBS 1X, AmericanBio, Natick MA) for 30 min at 37°C followed by 20 µl of Proteinase K (>600 U/ml, ThermoFisher Scientific, Waltham MA) and 5µl of RNase A / T1 (2 mg/ml / 5000 U/ml, Fisher Scientific, Waltham MA) for 5 min at room temperature. DNA was eluted in 35 µl of Nuclease-free water (Qiagen, Hilden Germany).

### RNA extraction from apple stigma

Deep frozen stigma samples were retrieved from -80 °C and 200 µl of 1X PBS (pH 7.4, Invitrogen, Carlsbad CA) + 0.001 % Silvet L77 (PlantMedia, Dublin OH) + beta mercaptoethanol (% vol/vol) were added to the tube. Epiphytic bacteria were detached by combining water bath sonication for 5 min and followed by full speed vortex for 30 sec as described in DNA extraction protocol. Stigma were removed from the tube and RNA was extracted using the RNeasy PowerLyzer Tissue&Cells Kit (Qiagen, Hilden, Germany) according to manufacturer’s instructions. Yield and quality were determined using QuBit and Agilent Bioanalyzer, respectively. Eluted RNA was depleted from ribosomal RNA using NEBNext rRNA Depletion Kit (Bacteria, New England Biolabs, MA) then used as template to generate cDNA using NEBNext Ultra RNA Library Prep Kit for Illumina (England Biolabs, Ipwich MA). The resulting cDNA assessed with a high-sensitivity Agilent 2100 bioanalyzer. Size profiles of cDNA fragments were consistent across samples.

### Profiling the microbiota of apple flower stigma

To study the apple flower microbiota, we followed previously described protocol(3, 18). Briefly, the V4 region of the 16S rRNA gene was amplified using primer set, 515F and 806R in triplicates from either isolated DNA or generated cDNA. PNA clams were added to PCR mixture to reduce the amplification of apple plastids and mitochondrial sequences(3). PCR product of triplicates were pooled purified, normalized using SequalPrep kit (Invitogen, Carlsbad, CA). Normalized amplicons were pooled together and submitted for sequencing using Miseq v2.2.0 platform (Illumina Inc, San Diego CA) at Yale Center for Genome Analysis.

### Sequences processing and community analysis

MiSeq forward and reverse reads were joined and demultiplexed using qiime2 pipeline (q2cli v2021.4.0)(54). PhiX and chimeric sequences were filtered out using qiime2-DADA2(55). Scripts used for sequences processing are available under https://github.com/hmamine/PPE/tree/main/reads_processing and raw sequencing reads are accessible at sequence read archive accession number PRJNA966219. Taxonomic classification of reads was performed using qiime2 feature classifier (q2cli v2021.4.0)(54) and on scikit-learn naive bayes trained classifier(56). Classifier was trained on 16S rRNA sequences (Green Genes 13.8.99) and the primer set 515F/806R. Reads were assigned to amplicon sequence variants (ASVs) with 100% sequence identity. To measure community alpha-diversity, Shannon and Observed ASVs were computed on unrarefied reads and using the R package “phyloseq”(57). To compute Faith’s phylogenetic diversity, the R package “picante”(58) was used on unrarefied bacterial reads. Count reads were normalized by cumulative sum scaling normalization factors(59) prior to computing Bray-Curtis distances and projecting constrained principal coordinate analysis and community homogeneity measured by distances to centroid. All R scripts used in this study are indicated at https://github.com/hmamine/PPE.

### *In vitro* mono-culture and co-culture growth assays

For mono-culture assay, cells from 5 ml of an overnight culture of *E. amylovora* strain 110, *Pseudomonas* CT-1059 or *Pantoea* CT-1039 were pelleted, washed, and re-suspended in equal volume of 0.5 × phosphate saline buffer (0.5× PBS). Cell concentration was adjusted to an initial OD_600_ of 0.08 in partial stigma mimicking media. Cells were incubated in 28ºC for 6 hours with shaking. For co-culture assays, bacterial cell suspensions were prepared similarly to mono-culture growth assay. Monocultures of each strain were adjusted to the same density, mixed in equal volumn to produce an initial OD_600_ of 0.08. Population of each strain and pH of the culture were determined after seven hours of growth. The colony forming units of each isolate was determined by plating on LB agar medium with selective antibiotics. *E. amylovora* and *Pseudomonas* CT-1059. were resistant to rifampicin and streptomycin, respectively. *Pantoea* CT-1039. was sensitive to both aforementioned antibiotics. Thus, *Pa* counts were determined by counting cells that were resistant to both antibiotics, and subtracting those counts from cells resistant to either antibiotic independently.

### RNA extraction *in vitro* co-culture assays

Bacterial cells were collected from mono- and co-cultures after 6h of growth at 28°C in partial stigma mimicking medium. RNA was extracted using the RNeasy PowerLyzer Tissue & Cells Kit (Qiagen) according to manufacturer’s instructions. The RNA was eluted in 50 µl nuclease-free water. Quantity and quality control of the RNA were assessed by using QuBit RNA Broad-Range Assay (Invitrogen) and with RNA ScreenTape analysis on Agilent TapeStation Bioanalyzer. Prior to Illumina library preparation, rRNA depletion was conducted using NEBNext rRNA Depletion Kit (Bacteria, New England Biolabs) according to the manufacturer’s recommended protocol. Then, DNA removal coupled with cDNA synthesis were conducted using NEBNext Ultra II RNA Library Prep Kit for Illumina (New England Biolabs) according to the manufacturer’s recommended protocol. Quantity check preformed with Qubit dsDNA HS Assay Kit Invitrogen). Two pooled samples were submitted for sequencing, each with 12 samples and a total concentration of 20 ng. Library Sequencing was conducted on an Illumina NovaSeq platform through services provided by the Yale Center for Genome Analysis (YCGA).

### Transcriptome analysis

Raw meta-transcriptome reads are accessible at sequence read archive accession number PRJNA966219. Adapter removal and sequences trimming were conducted using Trimmomatic(60). The full genome of each of *E amylovara* 110, *Pseudomonas* sp. strain CT-1059 and *Pantoea* sp strain CT-1039, available in the GenBank SRA under the BioProject accession number PRJNA693803(16), were used to build unique indexes for read mapping. Reads were quantified against the generated index using the alignment tool Salmon (v1.10)(61). Differential expressed genes were identified using DESeq2 R package(62) using normalized reads indicated by transcripts per million. Genomes were annotated using Prokka annotation tool (v.1.14)(63) and submitted to KEGG annotation database(22). All scripts are available at https://github.com/hmamine/PPE/tree/main/metatrans.

## Data availability

All sequencing reads used in this study are available at the NCBI Sequence Read Archive BioProject number PRJNA966219. All R scripts used to generate the figures could be accessed at https://github.com/hmamine/PPE.

## Acknowledgments

We thank Dr. Brian Kvitko (Univ. of Georgia) for providing miniTn7-pURR25DK-3xmCherry and miniTn7-pURR25DK-mTurquoise, Yale Center for Gen ome Analysis for providing support during sequencing and high-performance computing center at University of Connecticut. This study was funded by the USDA-NIFA-Organic Transitions grant 2017-51106-27001, USDA-NIFA-Agricultural Microbiome grant 2020-67013-31794, CAES Board of Control Research Award (2022)

, and USDA-Specialty Crop Block Grant (SCBG) through the Department of Agriculture, State of Connecticut.

## Supplementary materiel list

**Suppl. fig. 1. A**, box-plots indicate Faith’s phylogenetic diversity index of the flower microbiota in untreated, water-treated (mock) and *E. amylovora* (*Ea*) inoculated flowers. **B**, plot depicts constrained principal coordinate analysis of Bray-Curtis distances of the apple flower microbiota. Counts were normalized using cumulative sum scaling factor. Color indicates treatment and shape indicates flowers age (Day 2, Day 4) and days post-inoculation of *E. amylovora* (2dpi, 4dpi). Each circle indicates the microbiota composition of one individual flower, ten biological replicates were used for each treatment. Significant differences between groups were tested using Kruskal-Wallis test, ** indicates p-value < 0.01 and NS indicatesf a p-value > 0.05.

**Suppl. fig. 2. A** and **B**, box-plots depict Observed ASVs and Shannon index diversity metrics of the active (rRNA) and the total (rDNA) flower microbiota communities, respectively. Color indicates treatment and letter shows significant differences between groups. **C**, the plot shows principal coordinate analysis of Bray-Curtis (BC) distances of the apple flower microbiota using 16S rRNA (circle) and 16S rDNA (triangle) sequencing methods. Counts were normalized using cumulative sum scaling factor. Color indicates treatment. **D**, violin-plots show BC distances to water-treated (mock) samples of the active (rRNA) and the total (rDNA) microbiota communities. Differences between treatments were tested using Kruskal-Wallis followed by Conover’s multiple comparisons tests (p-value adjusted to Benjamini-Hochberg < 0.05). In graphs, mock indicates water-treated samples, *Ea* indicates *E. amylovora* inoculated samples and *Pa:Ps:Ea* indicates co-inoculation of *Pantoea, Pseudomonas* and *E. amylovora*, respectively.

**Suppl. fig. 3. A** and **B** show differentially expressed genes (DEGs) in *Pseudomonas* (*Ps*) upon co-culture with *Erwinia amylovora* (*Ea*) and *Pantoea* (*Pa*), respectively. **D** and **E** indicate DEGs in *Pantoea* upon co-culture with *Ea* and *Ps*, respectively. **G** and **H** depict DEGs in *E. amylovora* upon co-culture with *Pa* and *Ps*, respectively. Count data were fitted to negative binomial distribution and DEGs were determined based on the cutoffs of 1.5 of log2 fold change in relative expression and p-value < 0.05 (adjusted to Benjamini-Hochberg). Blue and red circles in the graphs indicate down- and up-regulated genes upon co-culture, respectively. Gray color in circles indicates genes that did not significantly change in expression. Numeric values indicate total count of genes up- or down-regulated in expression. C, F and I, venn charts indicate count of genes that were differentially expressed across different co-culture growth conditions in Ps, Pa and Ea, respectively. Left-panel graphs in (C), (F) and (I) indicates down-regulated genes in Ps, Pa and Ea, respectively. Whereas right-panel graphs (C), (F) and (I) indicates up-regulated genes in Ps, Pa and Ea, respectively.

**Suppl. fig. 4. A, B** and **C** show differentially expressed genes (DEGs) in *E. amylovora* (*Ea*), *Pantoea* (*Pa*) and *Pseudomonas* (*Ps*) belonging to the category of ABC transporters, prokaryotic defense systems and secretion systems, respectively. Count data were fitted to negative binomial distribution and DEGs were determined based on the cutoffs of 1.5 of log2 fold change in relative expression and p-value < 0.05 (adjusted to Benjamini-Hochberg). Blue and red colors in the heatmap indicate down- and up-regulated genes upon co-culture, respectively.

